# Resource presentation dictates genetic and phenotypic adaptation in yeast

**DOI:** 10.1101/2024.09.23.614544

**Authors:** Neetika Ahlawat, Anjali Mahilkar, Supreet Saini

## Abstract

Environments shape adaptive trajectories of populations, often leading to adaptive parallelism in identical, and divergence in different environments. However, how does the likelihood of these possibilities change with minute changes in the environment? In this study, we evolve *Saccharomyces cerevisiae* in environments which only differed in how sugar source is presented to the population. In one set of populations, carbon was presented as a mixture of glucose-galactose, and in the other, as melibiose, a glucose-galactose disaccharide. Since the two environments only differ in how the two monosaccharides are packaged, we call these environments „synonymous‟. Our results show that subtle changes in environments change the targets of selection between the two sets of evolved populations. However, despite different adaptive responses, pleiotropic effects of adaptation are largely predictable. Genome sequencing results demonstrate that small changes in the environment also strongly dictates the genetic basis of adaptation.

## Introduction

Environment dictates adaptive trajectories of populations^1–5^. Identical environments tend to drive adaptive parallelism^6–18^, while populations evolving in different environments often exhibit adaptive divergence^4,19–23^. The collateral effects of adaptation, in similar or dissimilar environments, have been reported to be wide-ranging and often predictable^24–26^.

However, are adaptation and its effects predictable when the evolution environments are “nearly-identical”? For instance, *Escherichia coli* populations evolved in either glycerol or lactate, exhibit phenotypic convergence despite differing in their gene expression profiles^8^. A recent study explicitly asks this question and examines *E. coli*‟s adaptation in „synonymous‟ environments. In this study, *E. coli* is evolved in environments that differ only in how glucose and galactose is presented – as a mix of the two monomers, lactose, or melibiose^27^. The findings of this study show that while adaptive responses of populations are non-identical, pleiotropic effects of adaptation, in a range of non-synonymous carbon environments, are largely predictable.

The above two examples show that adaptation in nearly identical environments can exhibit qualitatively different signatures. How does this variability in adaptation in nearly-identical environments change, as we change the taxonomic group of the evolving organism? Do the tools of predictability reported in the past still hold^27^?

To answer this question, we evolve *Saccharomyces cerevisiae* in synonymous environments – a glucose-galactose mix and melibiose^27^. Natural isolates or common laboratory strains of *S. cerevisiae* cannot hydrolyse lactose or melibiose. However, a *S. cerevisiae* strain from several decades ago, carries MEL1, an α-galactosidase responsible for extracellular hydrolysis of melibiose into glucose and galactose, and is capable of growth on melibiose as the sole carbon source^28^.

After evolution for ∼300 generations, we study the adaptive response, genetic basis of adaptation, and pleiotropic effects of evolution in the two „synonymous‟ environments. We show that adaptation in nearly-identical environments leads to non-identical phenotypic and genetic responses in the evolved populations. Despite differences in adaptive responses, pleiotropic effects of adaptation are predictable across non-synonymous environments. Our results provide insights into how even a slight change in the environment can lead to non-identical adaptive responses, a trend might be consistent across different taxa.

## Results

### Adaptation in nearly-identical environments is non-identical

We evolved six replicate populations of yeast in each of the two synonymous environments-a mixture of glucose-galactose, and melibiose. After ∼300 generations of evolution, we quantify growth rate (*r*) and yield (*K*) as measures of fitness in the evolved populations in their home environments.

We observe an increase in the fitness parameters (*r* and *K*) in all evolved lines relative to the ancestor (**Figure 1A**). Populations evolved in glucose-galactose mix, on average, exhibit higher relative increase in yield, compared to the populations evolved in melibiose (*one-tailed t-test, p value=0.0094*) (**Figure 1A**). On the other hand, populations evolved in melibiose exhibit a higher relative increase in growth rate, compared to the lines evolved in glucose-galactose (*one-tailed t-test, p value=0.0015*). These results indicate that a subtle change in the environment can change targets of selection.

**Figure 1.**
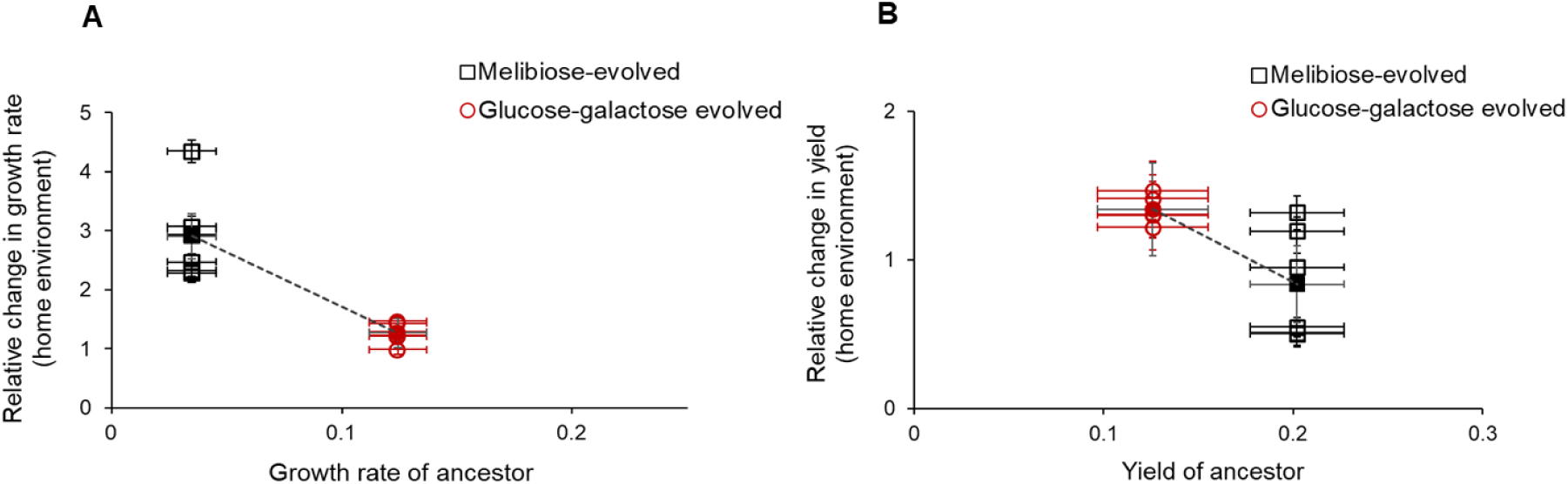
Populations exhibit different adaptive response in in nearly-identical environments. **(A)** Relative change in growth rate (***r***) and yield (***K***) in melibiose and glucose-galactose evolved populations is calculated with respect to the ancestor using, 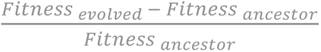. Filled symbols show the average relative fitness of the two sets of populations. Error bars represent standard deviation (SD) ± 1.96. **(B)** Range of fitness change within two sets of evolved populations is measured as the difference between maximum and minimum relative fitness across the replicate lines.

Next, to determine variability within replicate lines, we calculate the range of relative fitness gains as the difference between the maximum and minimum increases in the two parameters, *r* and *K*, in home environments. The six melibiose-evolved populations exhibit a wider range of the two fitness parameters as compared to glucose-galactose evolved populations (**Figure 1B**). Phenotypic variability within melibiose-evolved populations suggests that genetic targets of evolution are plenty in melibiose, relative to that in glucose-galactose.

### Pleiotropic effects in non-home synonymous environment

We quantify the fitness of populations in non-home synonymous environment by measuring the relative change in *r* and *K* of the two sets of evolved populations, with respect to the ancestor. Adaptation in the home environment leads to an increase in fitness in the non-home synonymous environments (**Figure 2A, 2B**).

**Figure 2.**
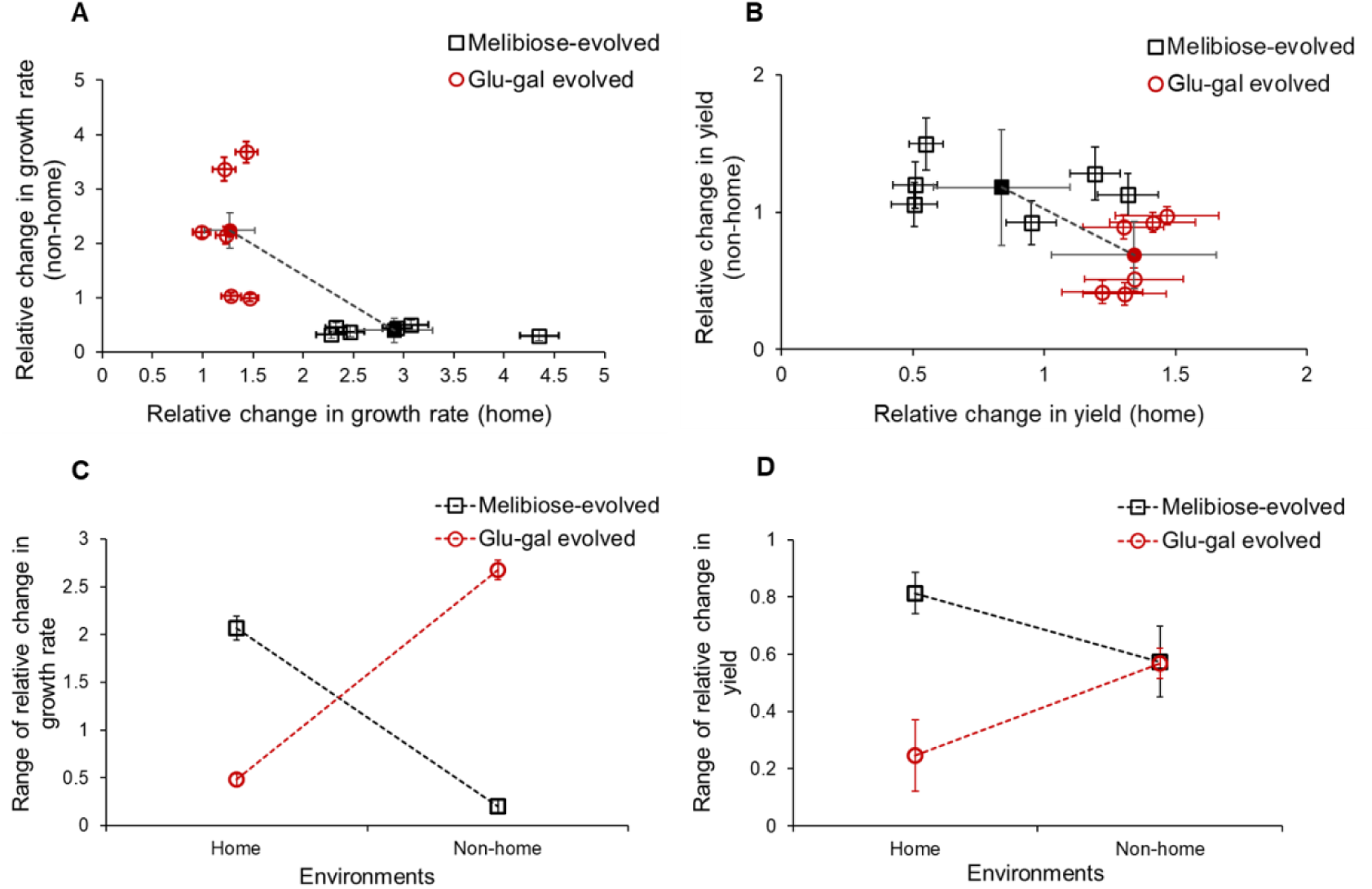
Pleiotropic responses of the two sets of populations in synonymous non-home environments. **(A)** and **(B)** fitness gains in home versus non-home synonymous environments. Filled symbols represent average gains in the relative fitness. Dotted line connecting average fitness values show the trend in fitness gains between melibiose and glucose-galactose evolved populations. **(C)** and **(D)** range of relative change in growth rate and yield across home versus non-home environments.

Even though the melibiose-evolved populations showed a greater increase in growth rate than the glucose-galactose evolved ones (**Figure 1A**), the average gains in growth rate in non-home environment was larger for the latter *(two-tailed t-test, melibiose-evolved p value=0.0006, glucose-galactose evolved p value=0.094)* (**Figure 2A**). Vice-versa, greater increase in biomass in glucose-galactose evolved populations (**Figure 1A**) shows smaller gains in biomass in non-home environment, relative to melibiose-evolved populations *(two-tailed t-test, p value for melibiose-evolved populations=0.121, p value for glucose-galactose evolved populations =0.0006)* (**Figure 2B**).

Next, we compare the variability in the fitness parameters (relative growth and relative yield) in home versus non-home environments **(Figure 2C, 2D)**. The variability in melibiose-evolved populations decreases in non-home environments, while that of glucose-galactose evolved populations goes up, for both the fitness measures under consideration.

### Pleiotropic effects in non-synonymous non-home environments

Next, we quantify fitness of the two sets of evolved populations in five non-synonymous sugar environments – (a) and (b) hexose sugars (monosaccharide): mannose and fructose, (c) neutral sugar: glycerol-lactate, (d) complex sugar (trisaccharide): raffinose, and (e) disaccharide: sucrose. We quantify relative growth rate (*r*) and yield (*K*) of the two sets of populations in non-synonymous environments, with respect to ancestor.

We compare the relative change in fitness of evolved populations in non-synonymous environments with that in home environments. We do not observe any correlated change between the relative change in growth rate in non-synonymous environments with the relative growth rate in their home environments *(melibiose-evolved, correlation coefficient (r)=0.16, p value=0.39; glucose-galactose evolved, correlation coefficient (r)=-0.047, p value=0.8)* **(Figure 3A)**. On the other hand, relative change in the yield in non-synonymous environments shows a negative correlation with the relative yield in home environments *(melibiose evolved r=-0.65, p value=<0.0001; glu-gal evolved r=-0.39, p value=0.033)* **(Figure 3B)**.

**Figure 3.**
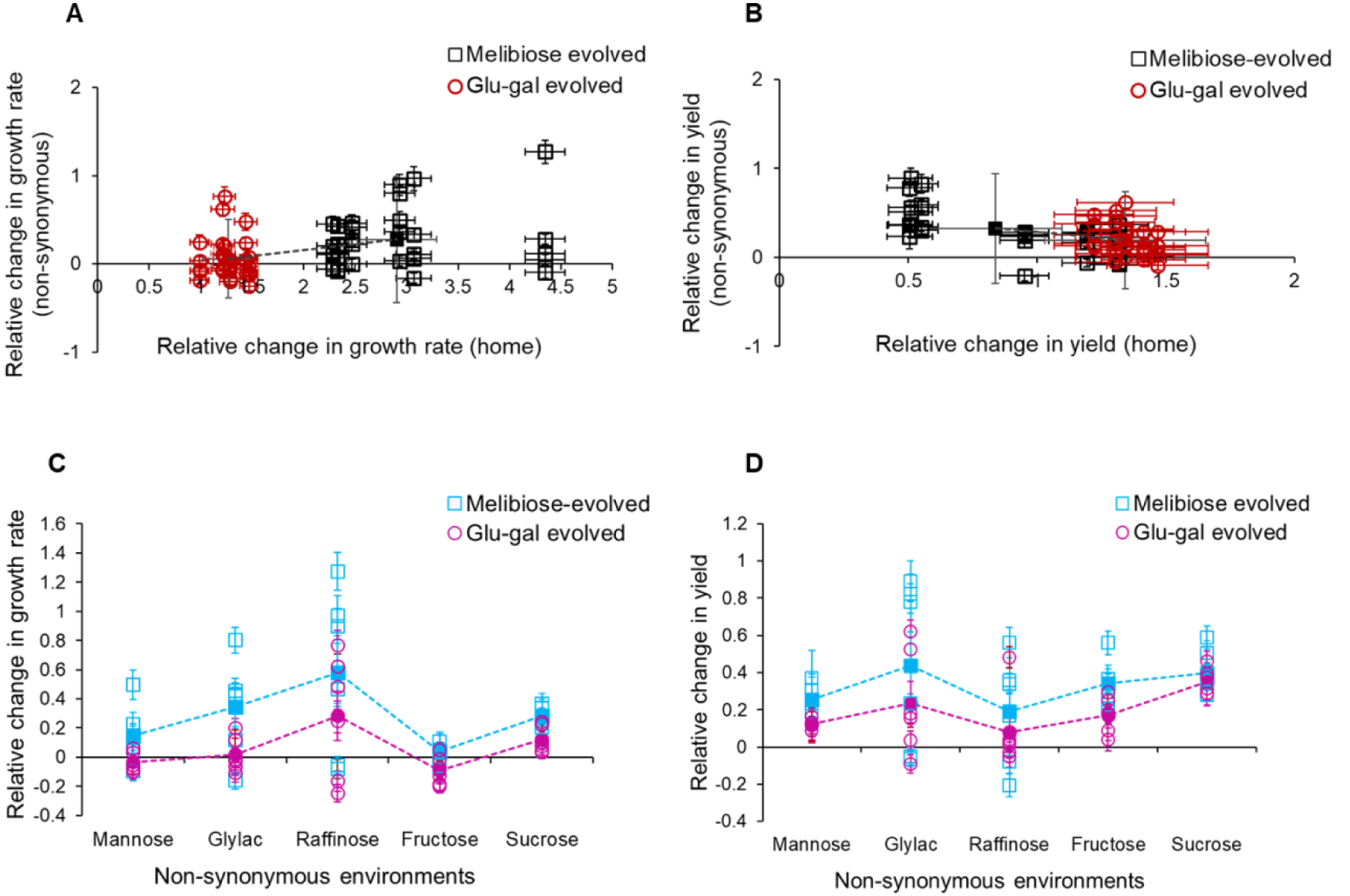
Pleiotropic responses of the two sets of populations in non-synonymous environments. **(A)** and **(B)** fitness gains in melibiose and glucose-galactose evolved populations in their home versus non-synonymous environments. Filled symbols represent average gains in the relative fitness. Dotted line connecting average fitness values show the trend in fitness gains between melibiose and glucose-galactose evolved populations. **(C)** and **(D)** fitness gains in the relative growth rate and yield in the two sets of evolved populations across all five non-synonymous environments. Mean values are represented with error bars indicating standard deviation.

Next, we compare the gains in the relative growth rate and yield between melibiose and glucose-galactose evolved populations across all five non-synonymous environments.

Although the two sets of evolved populations show positive pleiotropy, the analysis of variance results show a significant difference between the relative fitness of the two sets of populations, across non-synonymous environments (*for relative growth rate, F value = 10.49, p value = 0.002; for relative yield, F value = 5.72, p value = 0.02*) **(Figure 3C, 3D)**. Melibiose-evolved populations exhibit higher gains in the relative growth rate and yield across all five non-synonymous environments, as compared to glucose-galactose evolved populations **(Figure 3C, 3D)**. Pairwise comparisons between the two sets of populations show a significant difference between their relative growth rates across all five non-synonymous environments (**Table S1**). However, the difference between the relative yield of the melibiose and glucose-galactose evolved populations significantly differs only in the fructose and mannose environments (**Table S1, Figure 3D**).

Our results show that despite differences in the adaptive response of the two sets of evolved populations, their pleiotropic effects can be largely consistent across non-synonymous environments.

### Pleiotropic responses of populations can be predicted depending on ancestor’s fitness

We determine whether the relative change in fitness parameters (*r* and *K*) of the evolved populations across two categories of non-home environments (synonymous and non-synonymous), correlate with that of the ancestor‟s fitness.

In the two synonymous environments, relative change in growth rate and yield of the twelve evolved populations (melibiose evolved and glucose-galactose evolved) correlate negatively with the fitness of the ancestor *(growth rate r=-0.78, p=0.0026; yield r=-0.75, p=0.0048)* (**Figure 4A, 4B**). In case of non-synonymous environments, relative growth rate correlates negatively, whereas the relative yield correlates positively with the fitness of ancestor *(growth rate r=-0.48, p=0.0001; yield r=0.26, p=0.041)* **(Figure 4C, 4D)**. These results show smaller fitness gains in evolved populations, in environments where ancestor‟s fitness is higher, with an exception of biomass yield in non-synonymous environments.

**Figure 4.**
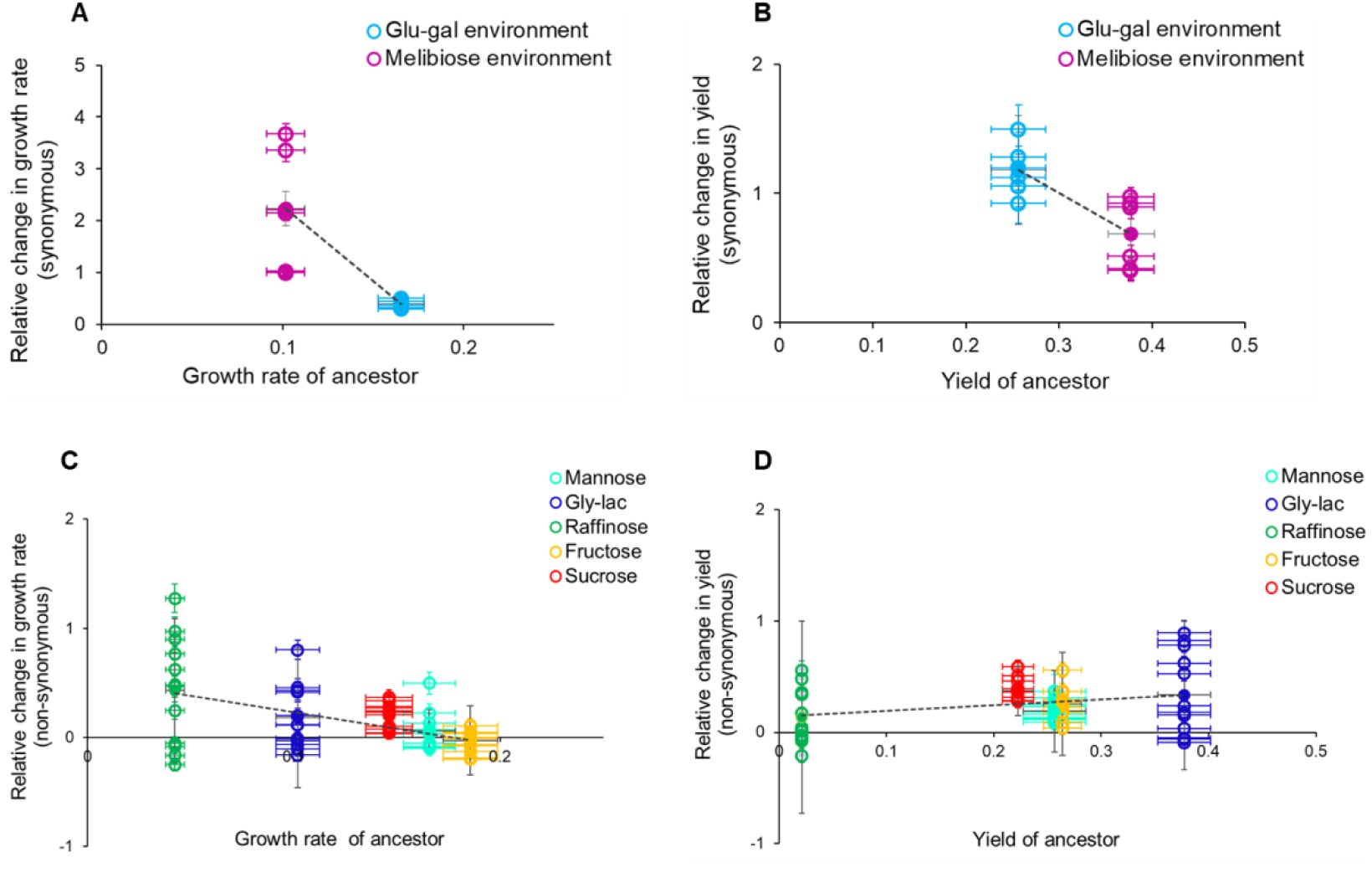
Predictability of pleiotropic responses in evolved populations based on ancestor’s fitness across synonymous and non-synonymous environments. A, B, C, and D show the comparison of relative changes in growth rate and yield between evolved populations and ancestor, across both synonymous and non-synonymous environments. Pleiotropic responses were predicted by measuring the fitness gains in twelve populations (evolved in melibiose and glucose-galactose) relative to the ancestor‟s fitness in different environments.

The observed patterns in **Figures 4A** and **4B** suggest that the ancestor‟s higher growth rate in glucose-galactose and greater biomass in melibiose correspond to lesser adaptation in those traits within their respective evolution environments (**Figure S1**).

These results suggest that provided the ancestor‟s fitness, adaptability of populations, and their pleiotropic responses can be predicted.

### Distinct genomic targets of adaptation in glucose-galactose and melibiose-evolved populations

To understand the genetic basis of adaptation in nearly-identical environments, we sequenced the complete genomes of populations evolved in either melibiose or glucose-galactose environments.

We observe a total of forty-one mutations in the two sets of populations. These mutations are spread over thirty-one genes (**Table 1 and S2**). Populations evolved in glucose-galactose had mutations in seventeen genes, and those evolved in melibiose had mutations in seventeen genes. Three genes were common targets between the two sets of populations (**Table 1**). Mutation in only one out of the seventeen genes is shared between two replicate populations evolved in glucose-galactose environment (**Table 1**). In contrast, mutations in five out of the seventeen genes are shared between at least two lines in the melibiose-evolved populations (**Table 1**).

**Table 1.**
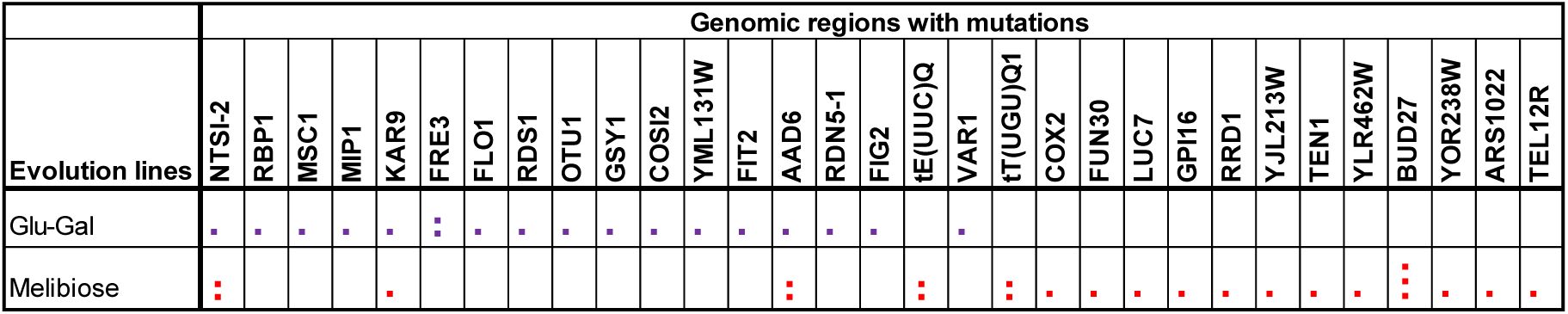
Genomic targets of adaptation. List of distinct targets of adaptation in the two sets of populations. Blue dots represent genes targeted in glucose-galactose evolved populations, and red dots show genes targeted in melibiose-evolved populations. Repeated dots in a cell represent mutational targets shared among replicate populations.

Distinct mutational targets within glucose-galactose evolved populations do not correlate with the phenotypic convergence of populations in their home environment. On the other hand, despite phenotypic divergence, some of the mutational targets are found to be shared within the melibiose-evolved populations.

These observations clearly demonstrate (a) non-identical targets of adaptation in nearly-identical environments, and (b) a lack of correlation between genotypic and phenotypic variability.

## Discussion

Adaptation in identical environments has been studied in the past^6–18,29^, and both parallelism and divergence have been observed depending on the evolutionary context. But how does the likelihood of these possibilities change with minute changes in the environment? How does this likelihood itself change with increasing complexity in the organism? We address these questions by evolving yeast populations in two nearly identical environments (glucose-galactose mix, and melibiose) that have been previously reported as „synonymous‟ sugar environments in an adaptive laboratory study conducted using *E. coli*^27^.

Our results, in concordance with the previous study using *E. coli*^27^, show non-identical adaptive responses in populations that evolved in „synonymous‟ environments. However, we observe that the targets of selection differ with the change in the organism - the biomass yield of melibiose-evolved yeast cells exhibit lesser fitness gains than their growth rate, unlike the melibiose-evolved *E. coli* cells where fitness gains were higher in biomass yield than in growth rate. *r-K* trade-offs have been reported in the past^30–34^, but the findings of this work show that evolution in a given environment leads to different trade-off patterns in different organisms. In addition, we also show that in *S. cerevisiae*, adaptation in an environment can be predicted based on ancestor‟s fitness.

At the molecular level, we see that the mutational targets are different for the two organisms in the same environment. While *E. coli* internalizes melibiose, yeast breaks it down into glucose and galactose outside the cell, and subsequently imports the monosaccharides. So, in both melibiose and glucose-galactose environments, importing the monosaccharides is vital for the yeast cells. Our expectation based on this mechanism of utilization was that the targets of selection would be largely shared between the glucose-galactose and melibiose-evolved populations. However, that is not the case. Only three genes were common targets. This suggest the role of different hexose transporters (HXT family)^35,36^ for glucose utilisation in the two synonymous environments.

Correlating the exact genotype with phenotype is challenging. As a result, predicting effects of adaptation based on macroscopic traits like fitness has been a recent focus^20,29,37–40^. In this context, our study shows that the ancestor‟s fitness serves as a predictor of pleiotropic effects of adaptation in non-home non-synonymous sugar environments. It remains to be tested if this rule of predictability holds in other environments as well.

Overall, this work explains the effects of minute environmental changes on adaptation and pleiotropic effects in eukaryotic asexual populations of yeast. However, the role of ploidy, which is an important factor in dictating evolutionary trajectories^41–44^ , has not been explored in this study. Also, most eukaryotes are sexually reproducing species, and we do not yet know how the effects of evolution in synonymous environments change if the evolving population reproduced sexually.

In yeast, some complex sugars such as, melibiose, sucrose, raffinose, are utilised via public-goods systems^45–47^. Like melibiose, these complex sugars are hydrolysed into monosaccharides in the extracellular environment, which are internalised in the cell. However, it remains unclear whether the „synonymous‟ sugar combinations such as, raffinose and a mixture of glucose-galactose-fructose; sucrose and a mixture of glucose-fructose, elicit identical adaptive responses. Studying its evolution in these synonymous environments would offer insights into how the presentation of resources influences the processes of diversification^48^, niche specialization^45,49–51^, and sympatric speciation^52–54^. Therefore, the findings of this work underscore the need to perform high-throughput experimental evolution using different model organisms in distinct sets of synonymous environments to elucidate the exact role of the evolution environment in dictating the likelihood and dynamics of different adaptive processes at the eco-evolutionary

## Material and Methods

### Strains used and media composition

We used haploid *S. cerevisiae* (Sc644 MATa/α MEL1 ade1)^45,55^ to setup selection experiments in the two „synonymous‟ environments. Evolution lines were evolved in yeast complete synthetic media (CSM) (composition: yeast nitrogen base, (NH_4_)_2_SO_4_, complete amino acid mix) containing either a mixture of glucose and galactose (0.1% each) or melibiose (0.2%).

To perform growth kinetic, we inoculated single colony of evolved populations (300^th^ generation) in YPD (1% yeast extract, 1% peptone and 2% D-glucose) in 25 × 150 mm borosilicate tubes, at 30°C on a rotary shaker at 250 rpm. We transferred 50μl of the overnight culture to 5ml fresh glycerol-lactate liquid media (3% glycerol, 2% of 40% lactate, complete AA mix and yeast nitrogen base). After 48 hours, cells were transferred through two rounds of subculture in 1:100 dilutions in 5ml glycerol-lactate media, each cycle lasting for about 48 hours.

### Evolution experiment

For the evolution experiment, Sc644 MATa and MATα were evolved either in glucose-galactose (0.1% each) or, in melibiose (0.2%) environments. In each sugar environment, there were six independent lines (3 MATa and 3 MATα) each derived from single ancestor clones of MATa and MATα strains. These lines were diluted 100x in 5ml yeast CSM and were serially propagated after every 24 hours. Evolution experiment was carried out in 25 × 150 mm borosilicate tubes under identical conditions i.e. 30℃ and at 250 rpm. These lines were evolved till ∼300 generations and were stored in 50% glycerol solution.

### Fitness measured in home and non-home synonymous environments

We quantified fitness of evolved strains in their home as well non-home synonymous environments. We performed growth kinetic for all six evolved lines across the two synonymous environments: (a) glucose-galactose, and (b) melibiose.

Home environment represents native/evolution environment, and non-home synonymous environment represents the non-native synonymous environment.

To perform growth kinetic, the desired volume of the glycerol-lactate yeast culture (mentioned above), was inoculated in 10ml of complete synthetic media containing either glucose-galactose or melibiose as the carbon source. Inoculum was measured such that all lines should start from a optical density (OD_600nm_) of 0.01. OD was recoded manually using Thermoscientific Multiscan Go plate reader. Yeast growth kinetic was conducted at 30℃ and OD was recorded at a time interval of 8 hours till 48 hours.

Growth rate was measured as described previously^56^. Maximum OD attained in each replicate population was used as a measure for the yield. For every single treatment, ODs were measured in triplicates in 96-well plate and the experiment was repeated three independent times.

### Fitness measured in non-synonymous environments

Growth kinetic was performed following the same protocol and under identical conditions as mentioned above, except that after the glycerol-lactate propagation, cells were inoculated to 10ml CSM containing different carbon sources (mannose, fructose, glycerol-lactate, sucrose, raffinose) at a concentration of 0.2%.

### Whole-genome sequencing

The evolved and ancestor populations were revived from freezer stock on a YPD plate. A single colony was inoculated and grown in 10 mL of liquid YPD for about 15-17 hours. The cell culture was harvested to isolate genomic DNA following the *S. cerevisiae* genomic DNA isolation protocol^57^. DNA concentration was measured immediately after DNA isolation using Nanodrop Spectrophotometer from Eppendorf (basic), and quality was checked by gel electrophoresis.

Genomic samples were sent for paired-end sequencing using Illumina NovaSeq 6000, with an average read-depth of 170bp.

Based on the quality report of fastq files, sequences were trimmed to retain only high-quality sequences for analysis and low-quality sequences were removed. The adapter trimmed reads were aligned to the respective reference genome, Sc288C, using Burrows Wheeler Aligner (BWA). Each sample had a minimum coverage of more than 30x. Variant calling was done for the samples using GATK and further annotated using SnpEff. Variants that were present in the ancestral strain were filtered out manually. After that, remaining SNPs were used for further analysis.

Raw sequencing data is available at https://www.ncbi.nlm.nih.gov/sra/PRJNA1150400.

### Statistical analysis

Pairwise comparisons between the mean values of the evolved populations were done using one-tailed/two-tailed t-test. For all the analysis, significance level was set to 0.05.

Relative changes in the growth rate and yield of the two sets of evolved populations were compared across home versus non-home environments, using pearson correlation.

## Acknowledgements.

This work was funded by a DBT/Wellcome Trust (India Alliance) grant (award no. IA/S/19/2/504632) to SS. NA was supported by IIT Bombay‟s Institute Postdoctoral Fellowship. AM was supported by the Council of Scientific and Industrial Research. We thank Pavithra Venkataraman for discussions.

