## Supplementary material for "Resource presentation dictates genetic and phenotypic adaptation in yeast"

### Resource presentation dictates genotypic and phenotypic adaptive trajectories in yeast.

**Supplementary file.**

#### Supplementary figure.

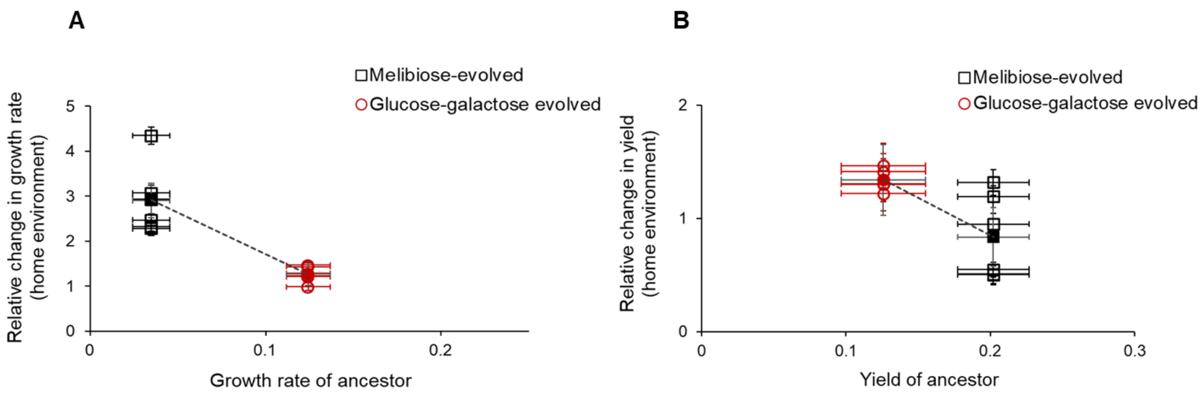

**Figure S1. Predictability of adaptation based on ancestor's fitness across synonymous environments.** Comparison between relative changes in growth rate and yield of evolved populations and ancestor, across their home/evolution environments. Adaptation in melibiose and glucose-galactose populations was predicted by comparing their relative fitness gains in home environment with that of ancestor's fitness in two synonymous environments.

**Supplementary tables.**

**Table S1.** Pairwise comparison (using t-test) between the relative change in the fitness of the two sets of populations across all five non-synonymous environments.

| **p-value table** | | | |
|---|---|---|---|
| Pairwise comparison between melibiose and glucose-galactose evolved populations | | | |
| **Relative growth rate** | | **Relative yield** | |
| **Environment** | **p-value** | **Environment** | **p-value** |
| Mannose | **0.027*** | Mannose | **0.0029**** |
| Gly lac | **0.017*** | Gly lac | 0.078 |
| Raffinose | **0.011*** | Raffinose | 0.124 |
| Fructose | **0.011*** | Fructose | **0.0008***** |
| Sucrose | **0.002**** | Sucrose | 0.240 |

**Table S2. Genomic targets of adaptation.** List of distinct targets of adaptation across 12 evolved populations.

| Evolution line | Genomic region | Type | Nucleotide base/AA change | Chromosome | Chromosomal position |
|---|---|---|---|---|---|
| Glu-gal1 | NTS1-2 | upstream deletion | | 12 | 460002 |
| Glu-gal2 | RBP1 | SNV | p.Ala862Val | 13 | 270018 |
| Glu-gal2 | MSC1 | SNV | p.Thr481Ala | 13 | 161620 |
| Glu-gal2 | MIP1 | SNV | p.Ser978Pro | 15 | 940454 |
| Glu-gal2 | KAR9 | SNV | p.Glu446Asp | 16 | 34350 |
| Glu-gal2 | FRE3 | upstream SNV | c.-2350T>C | 15 | 1053194 |
| Glu-gal3 | FLO1 | SNV | p.Thr19Ile | 1 | 203458 |
| Glu-gal3 | RDS1 | SNV | p.Arg29Trp | 3 | 311042 |
| Glu-gal3 | OTU1 | SNV | p.Leu115Pro | 6 | 45217 |
| Glu-gal3 | GSY1 | SNV | p.Thr18Ala | 6 | 176340 |
| Glu-gal3 | COS12 | upstream variant | c.-1378A>T | 7 | 1411 |
| Glu-gal3 | YML131W | upstream variant | c.-77C>T | 13 | 10120 |
| Glu-gal3 | FRE3 | upstream variant | c.-2350T>C | 15 | 1053194 |
| Glu-gal3 | FIT2 | upstream variant | c.-553G>C | 15 | 1058977 |
| Glu-gal4 | YFL056C- AAD6 | SNV | p.Ser51Asn | 8 | 15280 |
| Glu-gal5 | RDN5-1 | upstream deletion | n.-1984delC | 12 | 460415 |
| Glu-gal6 | FIG2 | SNV | p.Ala862Val | 3 | 270018 |
| Mel1 | tE(UUC)Q | upstream deletion | c.-4425delT | Mitochondrial genome | 30947 |
| Mel1 | tE(UUC)Q | SNV | c.-4422T>A | Mitochondrial genome | 30951 |
| Mel1 | VAR1 | SNV | p.Val71Asp | Mitochondrial genome | 49112 |
| Mel1 | tT(UGU)Q1 | deletion | c.-1215_-1214delAT | Mitochondrial genome | 62646 |
| Mel1 | COX2 | upstream_variant | c.-611T>A | Mitochondrial genome | 73147 |
| Mel2 | FUN30 | SNV | p.Lys386Glu | 1 | 116074 |
| Mel2 | LUC7 | SNV | p.Ser160Thr | 4 | 300521 |
| Mel2 | GPI16 | SNV | p.Ser148Ala | 8 | 483396 |
| Mel2 | RRD1 | SNV | p.Val156Ile | 9 | 55663 |
| Mel2 | YJL213W | SNV | p.Arg102Lys | 10 | 32467 |
| Mel2 | TEN1 | SNV | p.Lys55Glu | 12 | 167640 |
| Mel2 | NTS1-2 | deletion | n.-1571delT | 12 | 460002 |
| Mel2 | YLR462W | insertion | c.-4144_-4143insAC | 12 | 1065030 |
| Mel2 | KAR9 | SNV | p.Arg367Ile | 16 | 34112 |
| Mel3 | BUD27 | in-frame insertion | | 6 | 92409 |
| Mel3 | YOR238W | deletion | | 15 | 784052 |
| Mel3 | tT(UGU)Q1 | deletion | c.-1215_-1214delAT | Mitochondrial genome | 62646 |
| Mel4 | YFL056C-AAD6 | SNV | p.Ser51Asn | 8 | 15280 |
| Mel4 | BUD27 | in-frame insertion | | 6 | 92409 |
| Mel4 | ARS1022 | SNV | p.Pro207Ser | 10 | 715122 |
| Mel4 | NTS1-2 | upstream_variant | n.-1401G>A | 12 | 459833 |
| Mel5 | YFL056C | SNV | p.Ser51Asn | 8 | 15280 |
| Mel5 | YFL056C | SNV | p.Leu47Ile | 8 | 15293 |
| Mel6 | BUD27 | in-frame insertion | | 6 | 92409 |
| Mel6 | TEL12R | upstream variant | c.-4110T>C | 12 | 1064997 |
| Mel6 | TEL12R | insertion | c.-4144_-4143insAC | 12 | 1064997 |
